# Molecular surveillance of zoonotic bacterial pathogens in farm dwelling peridomestic rodents across the upper Midwest, USA

**DOI:** 10.1101/2021.07.28.454187

**Authors:** Nusrat A. Jahan, Laramie L. Lindsey, Evan J. Kipp, Bradley J. Heins, Amy M. Runck, Peter A. Larsen

**Author notes:** Corresponding author: Peter A. Larsen University of Minnesota 1971 Commonwealth Ave St. Paul, MN 55108.

## Abstract

The effective control of rodent populations on farms is a critical component of food-safety, as rodents are reservoirs and vectors for many foodborne pathogens in addition to several zoonotic pathogens. The functional role of rodents in the amplification and transmission of pathogens is likely underappreciated. Clear links have been identified between rodents and outbreaks of pathogens throughout Europe and Asia, however, comparatively little research has been devoted to studying this rodent-agricultural interface in the USA, particularly across the Midwest. Here, we address this existing knowledge gap by characterizing the metagenomic communities of rodent pests collected from Minnesota and Wisconsin food animal farms. We leveraged the Oxford Nanopore MinION sequencer to provide a rapid real-time survey of the putative zoonotic food- borne and other human pathogens. Rodents (mice and rats) were live trapped from three dairy and mixed animal farms. Tissues and fecal samples were collected from all rodents. DNA extraction was performed on 90 rodent colons along with 2 shrew colons included as outgroups in the study. Full-length 16S amplicon sequencing was performed with the MinION. Our data suggests the presence of putative foodborne pathogens including *Salmonella spp., Campylobacter spp*., *Staphylococcus aureus*, and *Clostridium spp.*, along with many important mastitis pathogens. A critically important observation is that we discovered these pathogens within all five species of rodents (*Microtus pennsylvanicus, Mus musculus, Peromyscus leucopus, Peromyscus maniculatus*, and *Rattus norvegicus*) and shrew (*Blarina brevicauda*) in varying abundances. Interestingly, we observed a higher abundance of enteric pathogens (e.g. S*almonella*) in shrew feces compared to the rodents analyzed in our study, however more data is required to establish that connection. Knowledge gained from our research efforts will directly inform and improve upon farm-level biosecurity efforts and public health interventions to reduce future outbreaks of foodborne and zoonotic disease.

## Introduction

Rodents are the largest taxonomic assemblage of mammals in the world and they are well known for harboring a plethora of zoonotic pathogens of concern for human and animal health [1]. Both native and invasive species of mice and rats have benefitted from human activities, especially agricultural systems. Rodents are a common hindrance of food production systems globally and they are known to transmit zoonotic pathogens to food animals and raw produce by contaminating the overall farm environment [2–5]. This transmission is largely due to the amplification of foodborne pathogens through the daily deposition of urine and fecal pellets into the production environment. For example, a single rodent within a barn or food-production facility can introduce upwards of 23 million *Salmonella* bacteria into production pipelines within 24 hours [6, 7]. However, the functional role that peridomestic (i.e., living in and around human habitations) rodents serve in the amplification and transmission of various zoonoses is likely underappreciated. For example, clear links have been identified between rodent pests and outbreaks of zoonotic diseases throughout Europe and Asia [8–12]; yet, comparatively little research has been devoted to studying this relationship in the United States [4, 13].

Specifically, regional studies focused on specific rodent species and their pathogen reservoir status across the diverse agricultural landscapes of the United States are lacking. Hence, our overarching research goal was to investigate the role of rodent pests on food animal farms as reservoirs or carriers of zoonotic pathogens, especially with respect to species-specific patterns.

Emerging genomic technologies are providing exciting new opportunities for the surveillance of zoonotic pathogens in diverse settings and environments. Next-generation sequencing platforms have opened the door to metagenomic profiling, particularly using the 16S rRNA gene. 16S rRNA gene sequence data is particularly useful as molecular marker for bacterial identification, including for pathogens with clinical relevance [14, 15]. The 16S rRNA gene has nine hypervariable regions (V1-V9) with varying levels, of which the V3 and V4 regions have been the most sequenced for host-associated microbiota and taxonomy assignation of bacteria [16]. Second-generation sequencing platforms, such as Illumina (e.g., MiSeq, HiSeq) yield very high quality, however the resulting sequence data consist of relatively short (∼300bp) reads, often permitting only the analysis of particular subregion of the full length (∼1,550bp) 16S rRNA gene [17].

Hence, the taxonomic assignment of reads and identification of bacterial taxa, specifically at the species level may be less reliable. On the other hand, the Oxford Nanopore Technologies (ONT) MinION sequencer is a third-generation sequencing platform based on single molecule synthesis technology with the capacity to sequence large DNA fragments to produce ultra-long sequencing reads (thousands to milions of bases in length)[18, 19]. For this reason, MinION sequencing technology can be used to sequence the full length 16S rRNA gene, thus, providing greater confidence in taxonomic assignments at the species level. This approach is important given that bacterial pathogenicity is typically considered a species or strain level phenomenon [20]. Although per-base accuracy of nanopore sequencing is lower (∼98%) than that of more commonly utilized next-generation sequencing platforms (e.g., Illumiina, PacBio) similar or even greater taxonomic resolution has still been achieved with this technology [21–24].

Furthermore, with continued product development, technological advances and better bioinformatic tools nanopore DNA sequencing will likely improve per base accuracy in the coming months and years [25, 26].

The The Oxford Nanopore MinION platform has been successfully applied in several studies including the characterization of bacterial mock communities [25,27,28]; microbiota profiling of species and tissues such as dog skin [29], canine feces [30], equine-gut [31], water buffalo milk [32], sea louse [33], microalgae [34]; identification of fungi [35], and plastic-associated species in the Mediterranean sea [36]. Additionally, metagenomic analyses of environmental samples obtained from glacier regions [37], aquatic environments (e.g., ocean water column [38], river water [39], wastewater [40], freshwater [41], building-dust [21], and the International Space Station [42], demonstrates the potential and applicability of nanopore sequencing for microorganism detection in diverse environments. Notably, nanopore sequencing has been applied to describe human gut [43], and nasal microbiota [44], along with the identification of the bacterial community associated with colorectal cancer tumors [45], and thrombus samples [46]. Furthermore, pathogen surveillance in EMS vehicles [15], analysis of infected prosthetic devices in real-time [47], *Salmonella* outbreak monitoring in hospitals [48], and detection of antibiotic resistance markers in clinical samples [49–52], shows the potential of the MinION platform as a pathogen surveillance tool. However, more studies are required to assess the plausibility of using this platform to analyze the bacterial community composition at the species level. To our knowledge there is only one previous study that analyzed rodent (mouse) fecal microbiota demonstrating the superiority of Nanopore sequencing at characterizing the bacterial community at the species level compared to the usage of short-read technologies for bacterial community characterization [22].

In the current study, we aim to describe the fecal metagenomic communities of farm-dwelling rodents and identify putative zoonotic pathogens potentially harbored by these rodents using the nanopore full length 16S amplicon sequencing method. Our study area included farms in the Upper Midwest of the United States (i.e., Minnesota and Wisconsin). Research focused on the rodent-farm interface in this geographic region is severely lacking, thus warranting study [1]. Our intent was twofold, to 1) better understand the farm-level rodent diversity in our study area and 2) to molecularly characterize rodent biological samples to assess their reservoir status of zoonoses of agricultural concern.

## Materials and Methods

### Rodent trapping and sample collection on farms

During the summer (2019) and fall (2019, 2020), we collected rodents from one dairy cattle farm (A), one mixed animal (dairy cattle and hog) farm (B) and another mixed animal (cattle and horse) farm (C). Farms A and B are located in Nicollet and Stevens counties of Minnesota, respectively and Farm C is in Sauk county of western Wisconsin. Farm A is a large dairy operation (∼20,000 dairy cattle), Farm B is a medium-sized operation (600 dairy cattle and 400 hogs), and Farm C is a small-sized family farm (mixed species, e.g. cattle and horses < 100 animals). Rodent activity was elevated on farms per observations by farm managers, particularly on Farm A where we observed hundreds of Norway rats (*Rattus norvegicus*) actively foraging around compost piles during daylight hours. All farms had poison bait stations, kill traps and cats as rodent control measures during the time of our visits. Four nights of rodent trapping were conducted at each study site using 150 Sherman live traps baited with oats.

Decontamination of all traps was performed using a 10% sodium hypochlorite solution (10 min soak) before and after each trapping event. All trapped animals were humanely euthanized following approved UMN IACUC protocols (protocol number 1809-36374A). Standard morphological techniques were used to identify rodents to species-level and metadata (e.g., species, age, weight, sex, body measurements) were collected for each individual animal. Biological samples (e.g., feces, intestine, colon) were collected and preserved (e.g., liquid nitrogen, freezer) for metagenomic analysis and further quantification of pathogens of interest. A schematic workflow of the overall study design is shown in Figure 1.

**Figure 1:**
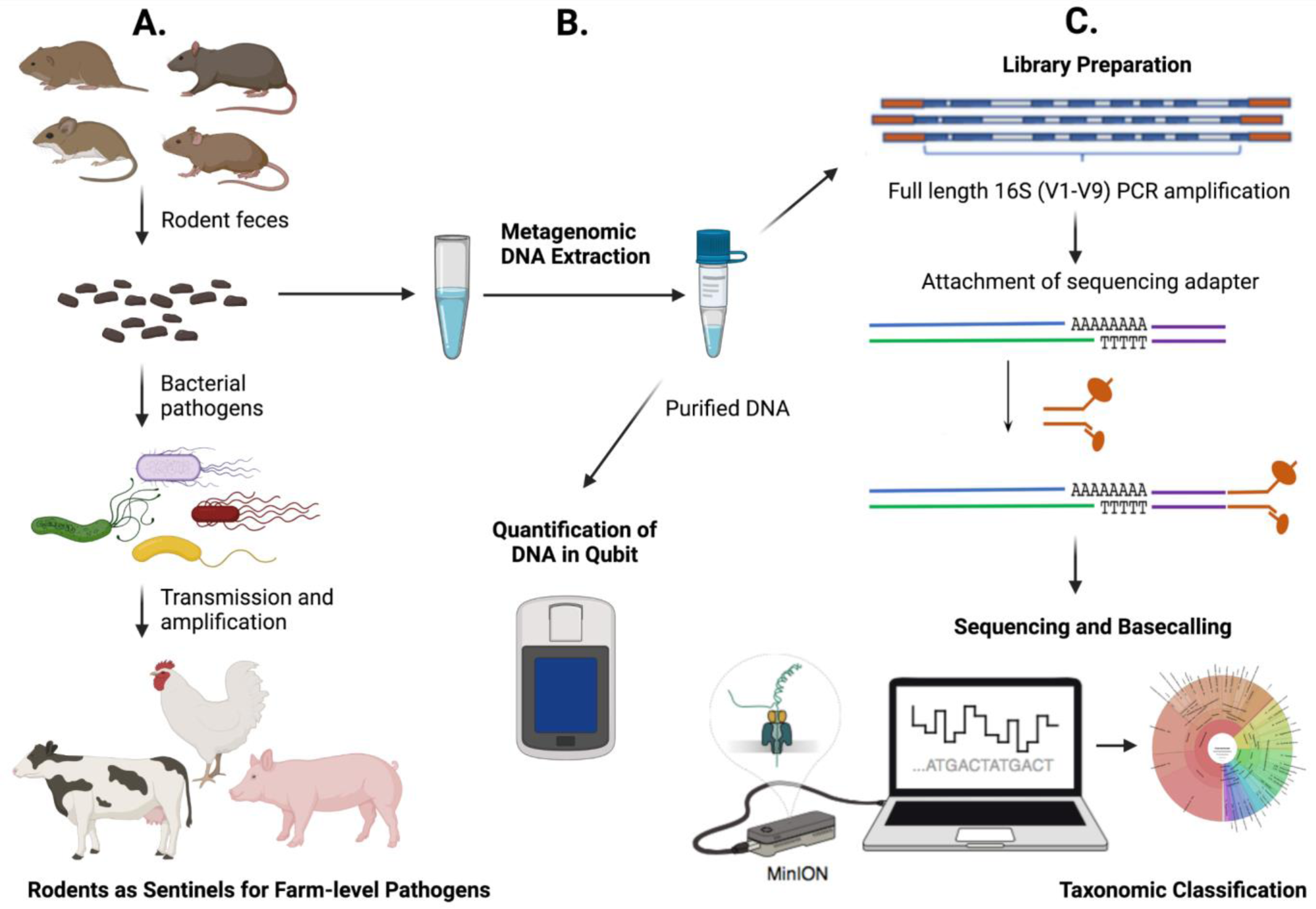
Workflow of the overall study design. **A:** showing rodents as amplifiers of bacterial pathogens through fecal deposition in the farm environment and possible transmission to farm animals. **B:** DNA extraction from rodent colon contents (i.e., feces) and quantification with Qubit fluorometer. **C:** Laboratory workflow to monitor bacterial communities from rodent fecal samples using Nanopore sequencing (Details in Materials and Methods).

### DNA extraction

Microbial metagenomic DNA was extracted with a QIAamp PowerFecal Pro DNA Kit (QIAGEN, Hilden, Germany). Snap-frozen rodent colon contents stored at -80°C were scraped and used for DNA extraction. Briefly, 250 mg of rodent colon contents were added to PowerBead Pro tubes and 800 μl of solution CD1 was mixed by vortexing. A bench top PowerLyzer 24 Homogenizer (QIAGEN, Hilden, Germany) was used for homogenizing the samples at 2000 rpm for 30 s, pausing for 30 s, then homogenizing again at 2000 rpm for 30 s to enhance cell lysis. PowerBead Pro tubes were centrifuged at 15,000 x g for 1 min and the resulting supernatant was transferred to clean microcentrifuge tubes. We used fully automated QIAcube connect instruments (QIAGEN, Hilden, Germany) for DNA extraction following manufacturer’s instructions. DNA concentrations were measured by fluorescence in a Qubit 4 fluorometer (ThermoScientific, USA) using the Qubit dsDNA BR Assay Kit (ThermoScientific, USA) following the manufacturer’s instructions.

### Nanopore library construction and sequencing

The 16S Barcoding Kit (SQK-RAB204; Oxford Nanopore Technologies, Oxford, UK) was used to prepare the amplicon library, following the manufacturer’s instructions for 1D sequencing strategy. The 16S region (1.5kb) of bacteria was amplified using specific primers (27F-1492R) and subsequently barcoded. This approach enables targeted sequencing of multiple samples and provides genus level resolution. Five sequencing runs were performed with a total of 70 samples, including 12 (run 1-4) and 22 (run 5) barcoded samples from individual rodents and shrews (colon contents). Briefly, genomic DNA samples were diluted to 100 ng/uL and amplification of the full-16S rRNA gene was performed by PCR with a reaction volume of 50 uL, using the primers 27F 5’- AGAGTTTGATCCTGGCTCAG-3’ and 1492R 5’-GGTTACCTTGTTACGACTT-3’, and Taq DNA polymerase LongAmp (NewEngland Biolabs, Ipswich, MA, USA).

Amplification was performed using Bio-Rad Laboratories PCR Thermal Cycler T100™ (Bio-Rad Laboratories, CA, USA) with the following PCR conditions: initial denaturation at 95 °C for 1 min, 25 cycles of 95 °C for 20 s, 55 °C for 30 s, and 65 °C for 2 min, followed by a final extension at 65 °C for 5 min.

The PCR products (50 μl each) were purified with 30 μl Agencourt AMPure XP beads and incubated in a HulaMixer for 5 min at room temperature. After the magnetic beads washing step, purified products were eluted in 10 uL of elution buffer (10 mM Tris-HCl pH8.0 with 50 mM NaCl). The amount and purity of the library was quantified using a Qubit 4 fluorometer (Thermoscientific, USA) following the manufacturer’s instructions. A single library was synthesized from DNA (100 or 50 fmol) of a pool of rodent colon contents. Libraries were pooled in multiplex mode following the addition of 1 μl of rapid adapter (Oxford Nanopore Technologies) and incubated at room temperature for 5 minutes. The amplicon library (11 μl) was diluted with a running buffer (35 μl) containing 3.5 μl of nuclease-free water and 25.5 μl of loading beads. Five nanopore sequencing libraries were separately run on FLO-MIN106 R9.4 (run 1, 2, 4, 5) and FLO- MIN111 R10.3 (run 3) flow cells (Oxford Nanopore Technologies) periodically after performing a platform quality control analysis. Sequencing runs were performed for 48 hrs. using the MinION control software, MinKNOW 4.0.20 (Oxford Nanopore Technologies).

### Bioinformatics analysis

After the completion of the sequencing run, raw signals in nanopore fast5 files were base-called (i.e. converting the electrical signals generated by a DNA or RNA strand passing through the nanopore into the corresponding base sequence of the strand) using Guppy (version3.2.2, Oxford Nanopore Technologies, UK), and a quality filter step was applied to retain only sequences with a mean Q-score ≥ 7. De-multiplexing of the barcoded samples was conducted using Porechop [53]. Adapter trimming and a second round of de-multiplexing were performed using Cutadapt 1.91 [54]. Only reads between 1,200bp and 1,800bp were selected for further analysis using Cutadapt, since the desired product size is ∼1,550 bp. Read statistics for each sequencing run were obtained using Nanostat and NanoPlot [55]. For taxonomic assignments Kraken2 [56] and Bracken [57] were used with the Greengenes (GG) database (https://benlangmead.github.io/aws-indexes/k2). While generating the Bracken classification report, a threshold of >100 reads were applied for higher confidence at the genus and species level. For visualization Krona tools and Pavian interactive applications were used to generate taxonomic charts and flow diagrams [58, 59]. The ggplot2 package (version 3.2.1) in RStudio software (version 3.3.3) was used to create a heatmap [60]. BioRender was used for illustrations and diagrams (Created with BioRender.com). Base-called data were uploaded to the EPI2ME interface, a platform for cloud-based analysis of MinION data, and WIMP (What’s in my pot) analysis was performed in parallel to compare results.

## Results

### Rodent trapping on farms

In total we trapped 90 rodents, 29 from Farm A, 43 from Farm B and 18 from Farm C (Table 1). From Farm C we also captured two shrews (*Blarina brevicauda)* and included those in the study as an outgroup. We identified five rodent species across our study sites, including three native (*Peromyscus maniculatus, P. leucopus*, and *Microtus pennsylvanicus*) and two invasive species (*Mus musculus* and *Rattus norvegicus*). The captured shrew species, *Blarina brevicauda,* is native to the Midwest. The majority of rodent captures were centered around feed bunks, grain storage sites, and within cow barns.

**Table 1:** Index of all captured animals from Farm A, B & C.

| **Rodent Species** | **Farm A** | **Farm B** | **Farm C** |
| --- | --- | --- | --- |
| House mouse (*Mus musculus*) | 27 | 21 | 0 |
| White-footed mouse (*Peromyscus leucopus*) | 0 | 13 | 11 |
| Deer mouse (*Peromyscus maniculatus*) | 1 | 4 | 0 |
| Meadow vole (*Microtus pennsylvanicus*) | 0 | 5 | 1 |
| Norway rat (*Rattus norvegicus*) | 1 | 0 | 6 |
| **Outgroup** |  |  |  |
| Shrew (*Blarina brevicauda*) | 0 | 0 | 2 |
| **Total = 92** | **29** | **43** | **20** |

### Nanopore sequencing workflow of full-length 16S rRNA for rodent microbiome analysis

We utilized the Oxford Nanopore MinION sequencer for rapid surveillance of rodent fecal microbiota. Full-length 16S rRNA (∼1,500bp) amplicon sequencing was conducted for metagenomic surveillance of foodborne zoonotic pathogens at the genus and species level. Colon contents were collected and used for DNA extraction (see details in the methods and materials section). We successfully generated full-length 16S amplicon sequencing data comprising more than 33 million DNA bases from 92 farm- caught rodent and shrew colon extracts. Run 1 and Run 3 included 24 rodent colon samples from the large conventional Farm (A), Run 2 and Run 4 included 24 samples from med-sized Farm (B), while Run 5 included all 20 samples from the small family Farm (C). Each of the sequencing runs included 12 molecularly barcoded samples except Run 5, which included 20 barcoded samples that were collected from individual rodent and shrew colons. The raw reads generated for the five runs from nanopore sequencing ranged from 3,531,607 to 9,445,273 with 500 to 509 average active pores and consisted of one dimensional (1D) forward, 1D reverse reads (Table 2). The mean quality score of the filtered reads ranged from 8.2 to 10.1 (Table 2), which indicated that the error rate of our sequencing was approximately 1 in 10 or 11 bases (∼10-11%), a result that is consistent with similar studies using nanopore sequencing [22, 61]. However, read depth of full-length 16S amplicons of 16X coverage, resulting in high-quality consensus, reduces concerns of per-base sequencing error [62]. To sort high-quality reads, the pass fast5 reads were sorted specifically from raw data using the Guppy base calling program. Although, Run 3 data (3,531,607 reads) showed fewer sequencing reads than other runs, the mean Q score (8.2), mean read length (1,625.40 bp) and read length N50 (1,593 bp) were comparable to the other runs (Table 2). For all the sequencing runs, the read length had a narrow length distribution, and the mean read length ranged from 1,132.70 to 1625.40 bp, which was close to the full-length of the 16s rRNA gene (about 1,550 bp).

**Table 2:** Statistics of nanopore sequencing (MinION) data.

| Run | Farm | Active Pores avg. | Read count | Mean read length | Mean Q score | Read length N50 | Total BP | QC > Q7 |
| --- | --- | --- | --- | --- | --- | --- | --- | --- |
| 1 | A | 509 | 5,011,566 | 1,132.70 | 9.3 | 1,564 | 5,676,494,189 | 85.80% |
| 2 | B | 500 | 4,460,963 | 1,178.30 | 8.3 | 1,587 | 5,256,288,918 | 85.50% |
| 3 | A | 509 | 3,531,607 | 1,625.40 | 8.2 | 1,593 | 5,740,290,501 | 68.50% |
| 4 | B | 500 | 9,445,273 | 1,477.10 | 10.1 | 1,576 | 13,952,018,041 | 85.40% |
| 5 | C | 504 | 5,829,408 | 1,472.30 | 8.4 | 1,564 | 8,582,667,914 | 77.70% |

However, for obtaining higher quality reads, a filtering program (Cutadapt) was applied to set the read length between 1200-1800 bp to discard any shorter or longer reads outside that specified range. After Guppy basecalling, demultiplexing, Porechop adapter trimming, and Cutadapt length trimming steps we filtered out around 20% of the initial raw reads from all runs. These polishing steps are necessary to ensure the presence of enough sequence similarity between the reads generated and precise matching against the 16S rRNA gene reference database. The longer reads that were removed during the filtering steps seemed to be the products of concatemers formed at the hairpin adapter ligation step, and the corresponding generation of long chimeric reads [22].

### Rodent fecal core microbiome and microbial diversity

The fecal microbiota composition was determined based on the nanopore sequencing data obtained with the taxonomy-supervised approach that allocates sequences directly into taxonomic bins based on their similarity. After the polishing step, all the long-read amplicons sequenced by Nanopore MinION were taxonomically assigned against the GreenGene (GG) reference (13_8 version) using Kraken2. After taxonomic classification, we obtained 96.25% of classified reads and 3.75% of unclassified reads. Total reads corresponding to Bacteria were at 96.25% and 0% reads were assigned to virus, fungi and protozoa. The microbial classifications were obtained at different taxonomic levels (e.g., division, phylum, class, order, family, genus, and species) for all of the 92 colon extract samples. Overall, the most abundant phylum for all five rodent species was Firmicutes (∼75% of total reads), followed in abundance by Bacteroidetes (12.5%), Proteobacteria (∼ 10%), and other phylum comprised of less than ∼ 2.5% from the total classified 80 phylum (Figure 2). To define microbial diversity and observe any specific patterns related to the different rodent species, we compared all five rodent species data at the genus and species level.

**Figure 2:**
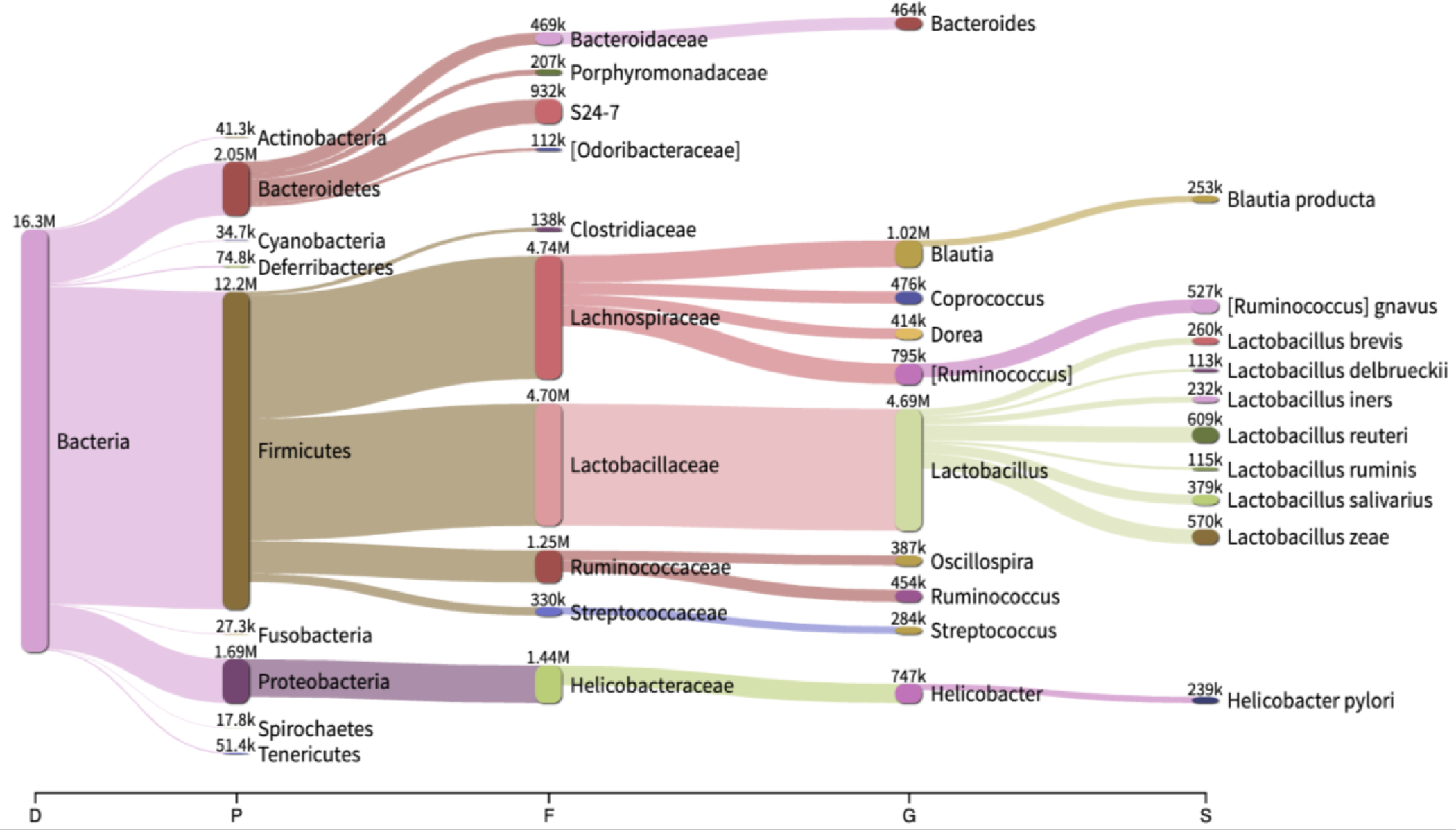
Core rodent fecal microbiome observed herein (n=90)

At the rodent species level, house mouse (*Mus musculus*) was the most captured species with a total of 48 animals caught from Farm A and B. After taxonomic classification of all the combined house mouse reads, 77 phyla, 886 genera, and 473 species were obtained. At the genus and species level, a filtering step was applied that included a threshold of 100 reads binned to the respective taxonomic level, which retained 261 genera and 181 species passing the threshold. For *M. musculus,* the most abundant genera were *Lactobacillus* (27.7%), *Ruminococcus* (10%), *Helicobacter* (8%), *Bacteroides* (7.6%), and *Blautia* (6.8%). Other abundant genus included fecal or mammalian gut microbiota (e.g., *Coprococcus, Faecalibacterium, Dorea, Roseburia, Oscillospira*) and potential human pathogens and bovine mastitis causing pathogens such as *Staphylococcus, Streptococcus, Bacillus*, and *Enterococcus*, each constituting more than 1% of the total classified reads.

The second most dominant rodent species was *Peromyscus* spp. with a total of 29 animals captured from Farm A, B, and C. After combining all *Peromyscus* reads and taxonomic classification, 58 phyla, 619 genera, and 344 species were obtained. After threshold filtering 139 genera and 89 species were retained. The most abundant genus was *Lactobacillus* (37.6%), followed in abundance by *Ruminococcus* (16.5%), *Blautia* (10.6.%), *Dorea* (4.6%), *Helicobacter* (4%), *Streptococcus* (3.4%), and other fecal related genera forming less than ∼ 3% of the total classified reads.

Brown rats (*Rattus norvegicus*) were captured from Farm A and C with a total of 7 animals in the category. Although we observed hundreds of *R. norvegicus* within Farm A, these rats are notoriously difficult to live trap using Sherman Traps during limited trapping durations. Future collections for this species will utilize a variety of trapping methods. All the combined nanopore reads were taxonomically classified into 47 phyla, 583 genera and 357 species. Threshold filtering retained 119 genera and 88 species, where *Lactobacillus* (22%), *Blautia* (17.3%), *Ruminococcus* (11.5%), *Streptococcus* (7.8%), and *Dorea* (6.5%) were most abundant. Other mammalian gut microbiota (e.g., *Oscillospira, Coprococcus, Faecalibacterium, Roseburia*) and potential human pathogenic genera (*Helicobacter, Prevotella, Staphylococcus*, *Clostridium, Bacteroides*) compiled more than 23% of the bacterial genera.

We captured a total of 6 meadow voles (*Microtus pennsylvanicus*) from Farm B and C. Taxonomic classification revealed 52 phyla, 542 genera and 302 species.

Subsequent threshold filtering retained 121 genera and 61 species for all combined reads. The most prominent genera in the meadow vole feces were *Lactobacillus* (24%), *Ruminococcus* (19.6%), *Blautia* (11.7%), *Oscillospira* (7.6%), and *Coprococcus* (5.3%).

While comparing species level composition, overall abundance of a few species were observed for all five rodent species (Figure 3). For example, most abundant *Lactobacillus* species included *L. reuteri, L. zeae, L. salivarius, L. delbrueckii, L. brevis, L. helveticus, L. ruminis,* and *L. iners.* Another dominant genus *Ruminococcus* represented *R. gnavus, R. torques, R. flavefaciens, R. bromii, and R. callidus. Blautia* species included *B. producta* and *B. obeum.* Whereas *Dorea formicigenerans, Roseburia faecis, Prevotella copri, Faecalibacterium prausnitzii, Oscillospira guilliermondii, Clostridium perfringens, Helicobacter pylori,* and *Coprococcus eutactus* represented single dominant species across all rodents. Furthermore, *Staphylococcus* species included *S. aureus, S. epidermidis, S. haemolyticus, S. sciuri* and *Streptococcus* species included *S. luteciae, S. anginosus, S. alactolyticus, S. infantis*.

**Figure 3:**
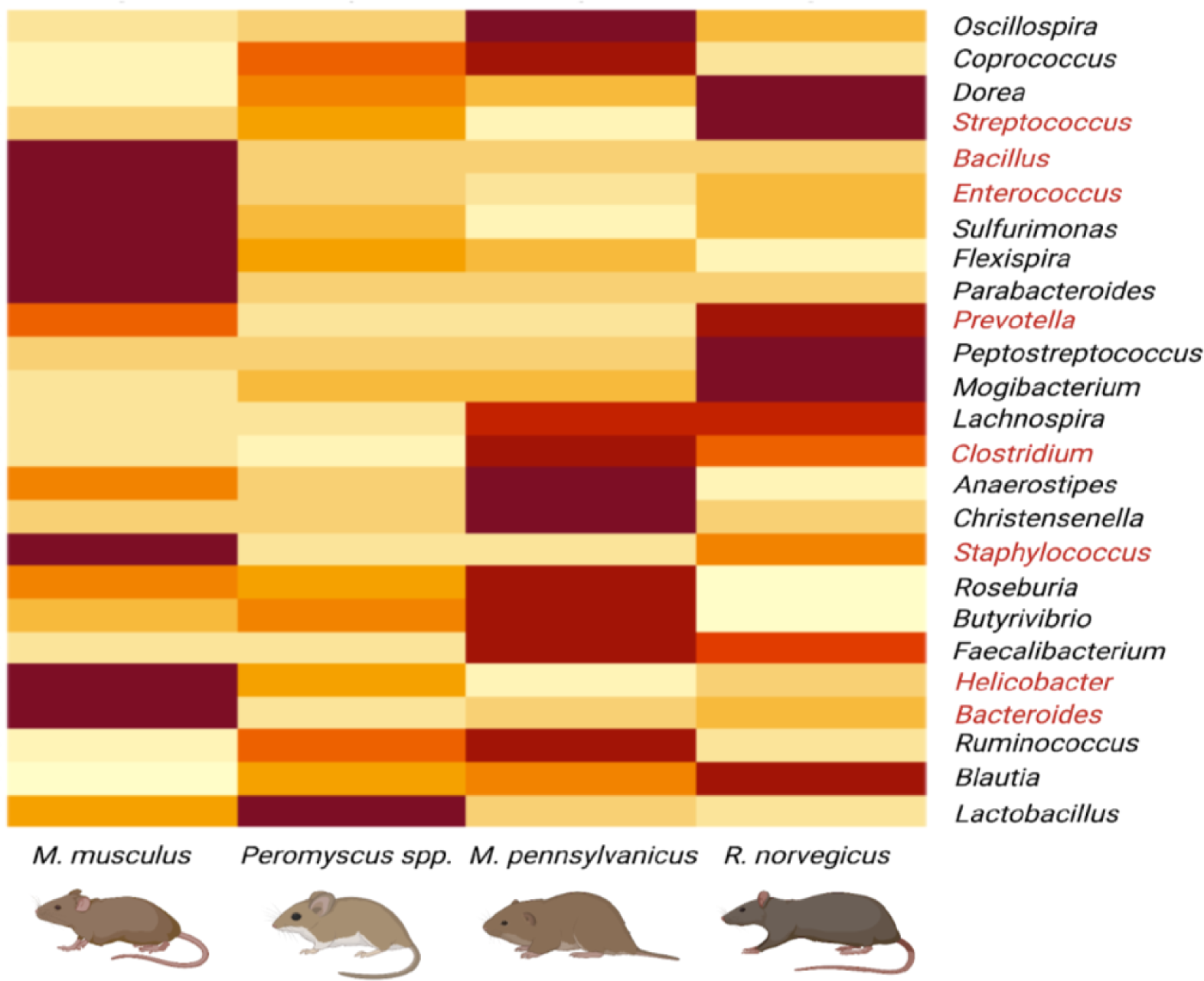
Heatmap of the most abundant (> 1%) bacterial genera identified by mapping 16S rRNA gene amplicons sequenced on Nanopore MinION against the GG reference database. Potential human pathogenic genera are labelled red in the legend.

### Shrew fecal core microbiome and microbial diversity

Two shrews from the same species *Blarina brevicauda* were captured from the small family farm (Farm C). Taxonomic classification analysis showed 28 phyla, 281 genera, and 178 species from the shrew sequencing reads. The most abundant phylum for both shrews was Proteobacteria (∼ 91% of total reads), followed in abundance by Firmicutes (8%), and other phyla comprised less than ∼ 1% from the total classified 28 phylum (Figure 4).

**Figure 4:**
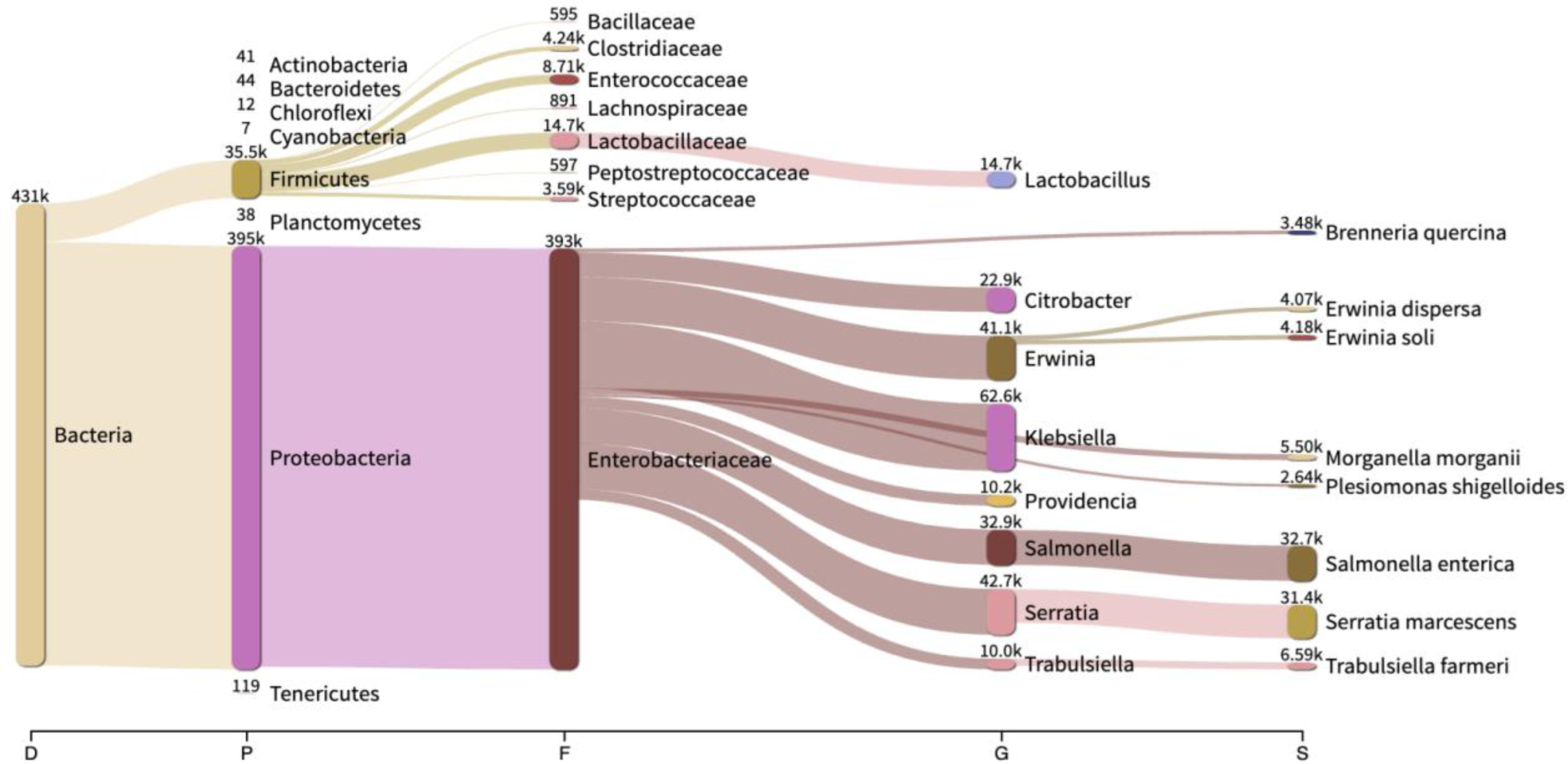
Shrew (*B. brevicauda*) fecal microbiome observed herein (n=2)

At the genus level, a total of 281 genera were identified and 36 genera retained after applying a threshold of 100 reads binned to each genus. The shrew fecal microbiome was rich in *Klebsiella* (18.8%), followed in abundance by *Salmonella* (16.8%), *Serratia* (15.7%), *Erwinia* (12.6%), and *Citrobacter* (6.2%). Moreover, it also contained other fecal-related and potential pathogenic genera including *Providencia, Enterococcus, Morganella, Yersinia, Enterobacter, Proteus, Clostridium, Plesiomonas, Vibrio, Bacillus, Pseudomonas, Staphylococcus,* and *Streptococcus,* representing less than 5% each of the total bacterial composition. The relative abundance of the 18 most abundant taxa determined at genus level is shown using bar graphs in Figure 5.

**Figure 5:**
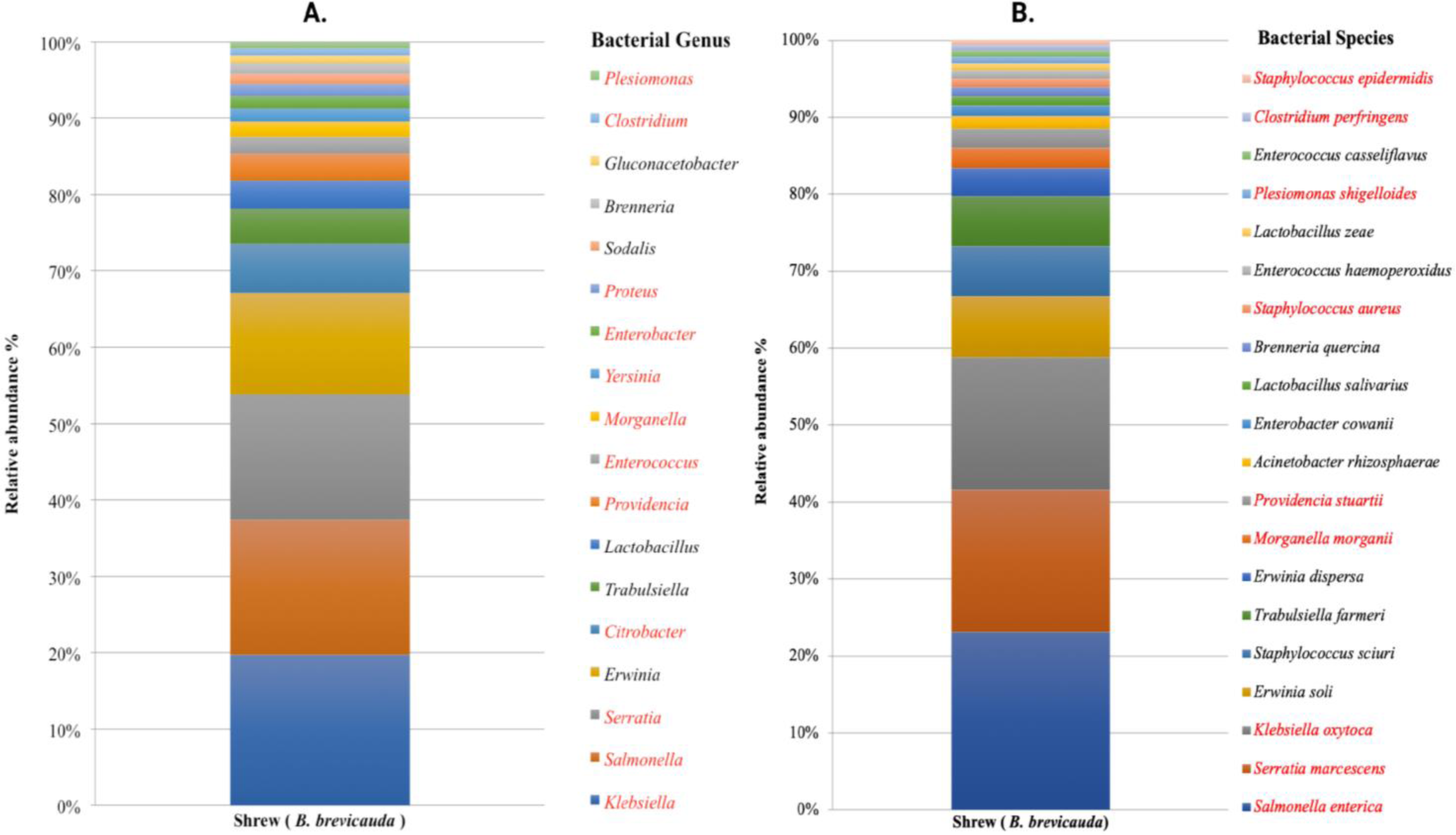
Shrew (*B. brevicauda*) microbiota representing > 1% relative abundance at the genus (A) and species (B) level.

Furthermore, analysis at the species level revealed a total of 178 species and 29 were retained after applying a threshold of 100 reads binned to each species. *Salmonella enterica* was the most abundant species with 21.5% reads assigned, followed in abundance by *Serratia marcescens* (17.3%), *Klebsiella oxytoca* (16%), *Erwinia soli* (7.4%), *Staphylococcus sciuri* (6.2%), and *Trabulsiella farmeri* (6%). Other abundant species included putative human and plant pathogens such as *Morganella morganii, Providencia stuartii, Enterobacter cowanii, Staphylococcus aureus,* and *Brenneria quercina,* each > 1% of the total bacterial species composition.

## Discussion

We utilized the MinION sequencing technology to characterize fecal microbial communities in peridomestic small mammals (i.e., rodents, shrews). DNA samples recovered from animal colon extracts were obtained from different dairy and mixed animal farms over a two-year period, and subjected to full-length 16S ribosomal RNA (rRNA) sequencing of amplicons for microbial identification. The 16S rRNA gene has been shown previously to be a useful molecular marker for bacterial identification, including for pathogens with clinical relevance [14]. This sequencing method and analysis pipeline demonstrates the convenience of this technology being highly portable, relatively inexpensive, small, and scalable for the rapid surveillance of pathogenic organisms in peridomestic pests [63–67]. Additionally, we demonstrate the ability to identify important rodent species in the dairy farms to monitor, inform biosecurity practices and suggest the potential role of rodent pests on the spread of pathogenic microorganisms and describe the rodent microbiome in general.

### Substantial compositional differences between animal species microbiome

To the best of our knowledge, this is the first study to reveal and compare the composition of bacterial communities in gastrointestinal tracts of wild caught rodents (e.g., *M. musculus, Peromyscus spp., M. pennsylvanicus, R. norvegicus*) using the third-generation sequencing technology. Moreover, our study is the very first published study revealing the gut microbiota of the Northern short-tailed shrew (*B. brevicauda*). Rodents are considered to be the most commensal synanthropic animals living in close proximity to humans and feeding on/foraging our food systems. Whereas, shrews (*B. brevicauda*) naturally live in woodlands, cultivated fields, vegetable gardens and mainly feed on invertebrates [68, 69]. However, they often retreat into barns, cellars and sheds during winter, which provides more opportunities for human–animal contact, and therefore higher rates of bacterial transmission are likely [70]. Therefore, commensalism and food- habit (omnivore vs. insectivore) might explain the difference in gut microbial structure between rodents and shrews. In our study, we observed that the bacterial communities of rodents and shrews (insectivore) were considerably different, even though they were captured from the same habitats. However, all the rodent species from different habitats (Farm A, B and C) had similar core microbiota indicating host tropism, also reported by a previous study investigating gastrointestinal helminths diversity of free ranging Asian house shrews [70]. Hence, the difference between the animal species (i.e., rodents and shrews) was greater than the difference within species (i.e., all rodents).

For all five rodent species, phylum level gut microbiome was similar to that of the human gut microbiome with Firmicutes, Bacteroidetes and Proteobacteria comprising more than 97% of the gut microbiota [71, 72]. Firmicutes was the most abundant phylum in all rodent species, ranging from 64% to 91.5%. In shrews, the most abundant phylum was Proteobacteria, covering 91.7% of the total shrew microbiota. This suggests a greater similarity in core gut microbial composition between rodents and humans than between shrews and humans. At the genus level composition, *Lactobacillus* was the top genus for all the rodent species in our study, which is in line with the findings of a previously reported study on laboratory rodent microbiomes [73]. This finding implies that the core fecal bacterial composition of wild and laboratory rodents share a degree of similarity.

Additionally, *Lactobacillus* constitutes a significant component of the human gut microbiome as well [74], reiterating the similarity between the composition of gut bacteria in humans and rodents. Furthermore, *Ruminococcus* was the second most prominent genus in all mouse species (*M. musculus, Peromyscus spp., M. pennsylvanicus*), whereas *Blautia* was prominent in rats (*R. norvegicus*). On the other hand, *Klebsiella* was the most prominent genus in shrew feces followed by *Salmonella*, *Serratia* and *Erwinia*. Whereas, *Clostridium* was the most abundant genus in the Asian house shrew (*Suncus murinus*) [75]. This might be explained by geographical location, habitat, feed, shrew species variation and limited sample size.

### Potential foodborne, bovine mastitis and other pathogens

We identified a higher relative abundance of potential human and animal pathogens in shrew fecal samples than in rodent fecal samples. In brief, the resulting data indicate the presence of multiple putative foodborne pathogens from the Enterobacteriales order including *Salmonella* spp., *Shigella* spp., *Plesiomonas shigelloides*, *Yersinia* spp., and *Escherichia coli*, in all small mammal species (Figure 6). Other top foodborne pathogens observed in our analysis included *Listeria monocytogenes*, *Campylobacter* spp., *Clostridium perfringens*, *Vibrio* spp., and *Staphylococcus aureus*. These are mentioned as major bacterial pathogens that cause foodborne illness and hospitalization in the USA and all over the world each year [76].

**Figure 6:**
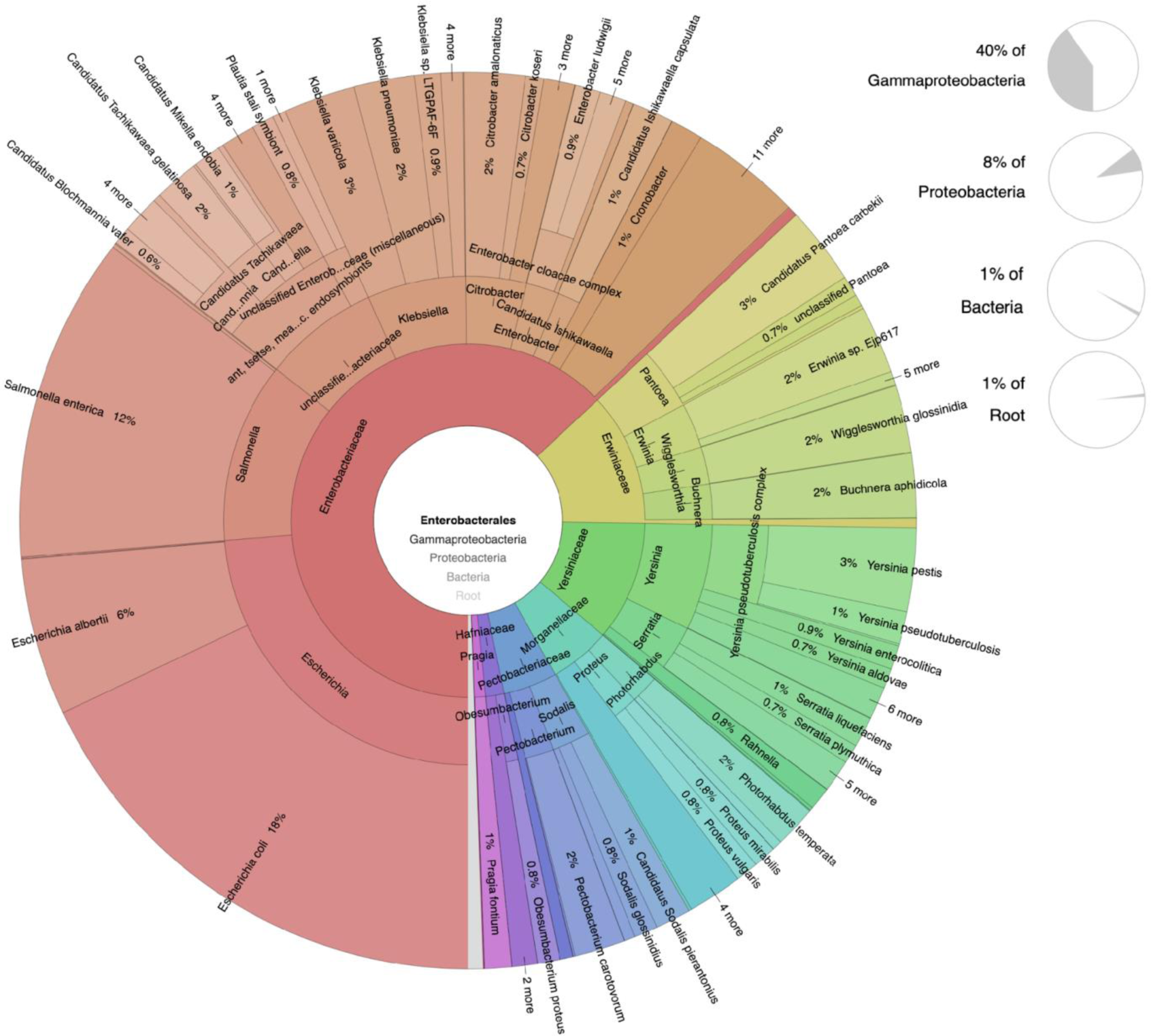
Krona plot showing overall (n=92) bacterial species abundance in the Enterobacteriales order.

Further, each of these bacterial foodborne pathogens were shown to be carried by different native and invasive rodent species all over the world [1].

Interestingly, *Salmonella enterica* was the most abundant species (∼21.5%) in the shrew feces and much lower abundance was observed for the rodents (Figure 7). On the contrary, *Clostridium perfringens*, *Bacillus* spp., and *Staphylococcus aureus* seemed to be carried by all rodents in higher abundance compared to shrews. Additionally, *Vibrio* spp. and *Campylobacter* spp. were observed in comparatively higher abundance in *R. norvegicus* and *M. musculus,* respectively. These bacteria can be transmitted by contaminated food and water, and they had a greater relative abundance in fecal samples of shrews than in those of rodents. These observations indicate that *B. brevicauda* may pose a high potential risk for spreading enteric pathogens via water or food contaminated with feces. Furthermore, farm workers can easily get infected by these pathogens carried by the small mammals through direct and indirect contact with fecal droppings.

**Figure 7:**
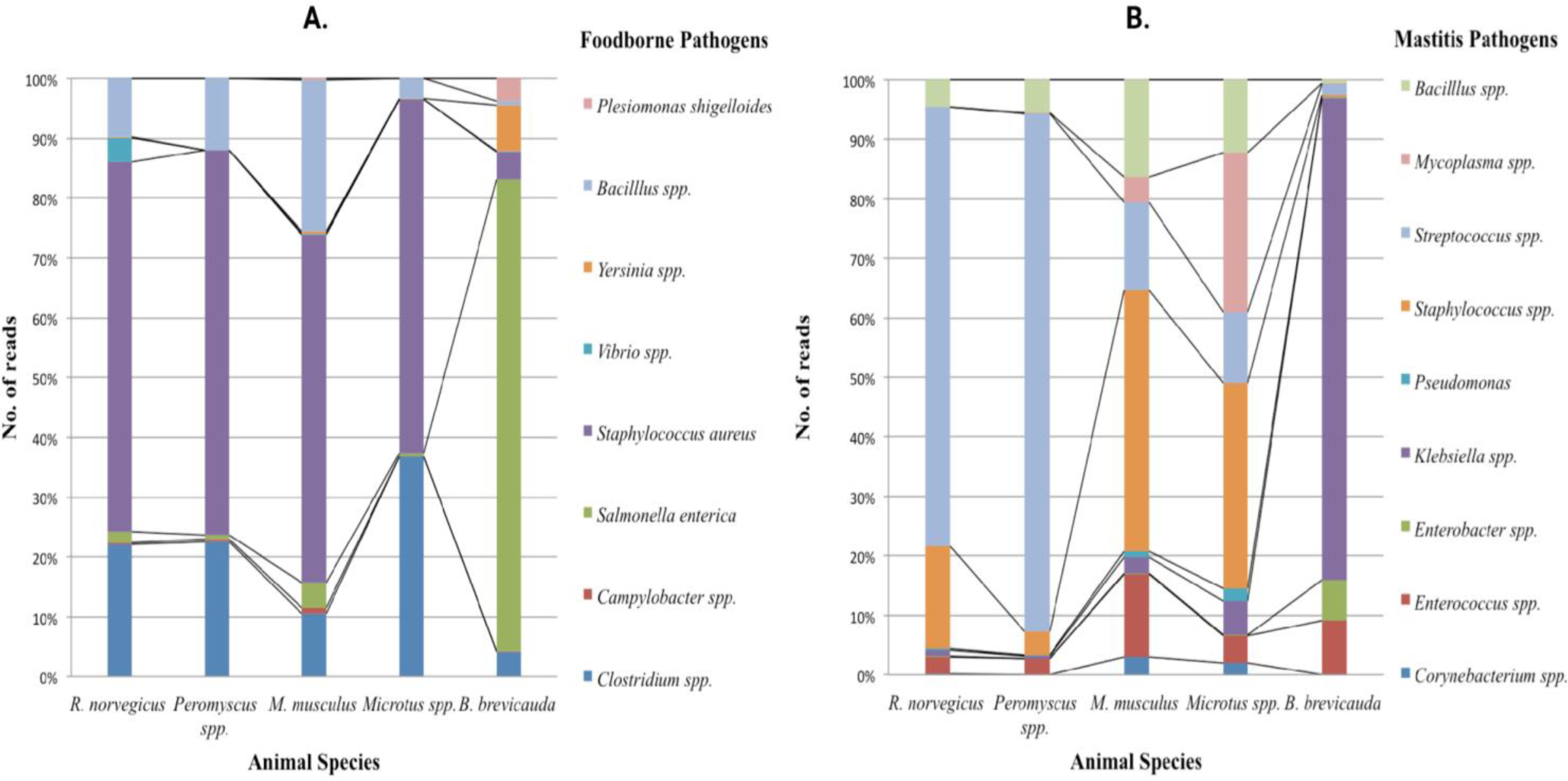
Abundance of Foodborne (A) and Mastitis (B) pathogens in all small mammal species.

Mastitis is a leading cause of cow culling and causes a great economic loss to the dairy industry [77]. Since a majority of the samples were collected from a large-sized (A) and medium-sized (B) dairy farm, we investigated the potential presence and abundance of pathogens causing bovine mastitis in the fecal samples of the farm dwelling rodents. Many important mastitis-causing pathogens including *Streptococcus* spp., *Staphylococcus* spp., *Klebsiella* spp., *Enterococcus* spp., *Enterobacter* spp., *Mycoplasma* spp., and *Corynebacterium* spp. were observed in varying abundance in the six species of small mammals (rodents and shrews) collected from each farm. *E. coli, Streptococcus* spp., *Staphylococcus* spp., and *Corynebacterium* spp. are well known environmental mastitis pathogens [78], and the presence of these pathogens in the resident rodent population of each dairy farm is a putative health risk to the resident cattle. Our metagenomic data indicate that the rodents sampled during our trapping events are possible reservoirs of mastitis pathogens and have the potential to continuously introduce these pathogens into the dairy farm environments. Moreover, they can amplify and mechanically vector these pathogens from sick to susceptible animals through pathogen amplification as described in [1]. Additionally, while comparing the relative abundance of mastitis pathogens from rodent colons from Farm A, B, and C (Fig 7B), the number of reads from house mouse (*M. musculus*) is higher compared to other rodent species for most mastitis pathogens, despite a similar depth of sequencing coverage. We hypothesize that because house mice cohabitate with cows inside the barns, they are exposed to a greater overall amount of mastitis pathogens by direct and indirect interactions with the large dairy herd.

Bacterial pathogens detected in fecal samples of rodents and shrews can cause human diseases by various routes, including saliva, urine, feces and contaminated water and food. In our study, fecal samples from all rodents also contained a variety of human pathogens in high abundance including *Helicobacter pylori* that causes chronic gastritis, gastric ulcers, and stomach cancer [79]; *Prevotella copri* that is associated with the pathogenesis of Rheumatoid Arthritis [80]; pathogens related to nosocomial infections such as *Morganella morganii*, *Serratia marcescens, Providencia stuartii* [81, 82]; and opportunistic pathogens associated with splenic abscess (*Parabacteroides distasonis*) and anaerobic peritoneal infections (*Bacteroides fragilis*) [83, 84]. Additionally, rodents are the known reservoirs of some severe infectious zoonotic diseases, such as bartonellosis, leptospirosis, plague, and rat-bite fever [1]. However, in our study, none of the fecal samples were positive for *Bartonella, Leptospira, Yersinia pestis*, *Streptobacillus moniliformis* and *Spirillum minus*. Similar findings were reported by He et al., 2020 investigating fecal and throat microbiota of Norway rats (*R. norvegicus*) and Asian house shrews (*S. murinus*) [75]. Hence, results from our study indicate that these infectious pathogens might not be shed in feces of the rodents, thus not detected in our study because of the sample type.

Both rodents and shrews are commensal animals that live and feed in closer proximity to human populations than most other mammals and may serve as potential sources of infectious zoonotic diseases to humans via pathogen amplification and cross-species transmission. However, less attention has been paid to these small mammals inhabiting our food processing system including food animal farms, fresh produce lands, and processing facilities. In our study, more sequences from the Northern short tailed shrew were annotated as potential foodborne pathogens than those from rodents. Therefore, *B. brevicauda* might be a more important reservoir of bacterial foodborne pathogens than rodents, suggesting that more attention should be paid to *B. brevicauda* in the prevention of foodborne zoonotic diseases in the future. However, more rigorous studies with larger sample sizes focusing on small mammal’s gut microbiota, specifically on *B. brevicauda* are required to support and confirm these findings.

## Limitations

Rodent trapping is unpredictable and dependent upon local conditions; hence it is difficult to determine species diversity. Additionally, we were not as successful trapping rats (*R. norvegicus*) compared to mice, despite visual observation of their abundant presence and activity in Farm A. Enhanced trapping efforts with pre-baiting measures to acclimate the rats and use of repeater live traps instead of Sherman traps might prove as useful strategies to overcome trap avoidance behaviors exhibited by *R. norvegicus* [85]. Nevertheless, our personal observation during trapping and subsequent results from data collected indicate that rodents are a problem in any agricultural setup and are potential reservoirs of many putative zoonotic foodborne and mastitis-associated pathogens.

The long read length, depth of coverage, and rapid results make the Nanopore sequencing method attractive for a molecular surveillance tool. Although Nanopore sequencing has a higher per-base accuracy error rate than other next-generation sequencing methods [61], we are basing our initial taxonomic classification on read depths of over 30 million full-length 16S (1.5 kb) reads. Moreover, the flow cell chemistry for the Oxford Nanopore MinION sequencer is constantly being improved upon and we used the latest R 9.4 and R10 Flow Cell sequencing technology that has improved per base accuracy compared to earlier flow cell versions (www.nanoporetech.com). However, a high number of sequencing reads for putative pathogens does not necessarily indicate the absolute presence of the organism. Standard culturing techniques and molecular methods are still required to confirm the presence of putative pathogens. Nevertheless, for our purpose of a primary screening tool, this cost effective and rapid method achieved our surveillance goals.

## Acknowledgements

The authors thank the University of Minnesota Agricultural Research, Education, Extension and Technology Transfer program for startup funds awarded to PAL. Bioinformatics were supported using tools available from the Minnesota Supercom- puting Institute.

